# To composite or replicate: how sampling method and protocol differences alter stream bioassessment metrics

**DOI:** 10.1101/847327

**Authors:** Lusha Tronstad, Oliver Wilmot, Darren Thornbrugh, Scott Hotaling

## Abstract

Aquatic invertebrates are excellent indicators of ecosystem quality; however, choosing a sampling method can be difficult. Each method and associated protocol has advantages and disadvantages, and finding the approach that minimizes biases yet fulfills management objectives is crucial. To test the effects of both sampling methods and sample handling – i.e., to composite samples or leave them as replicates – we collected aquatic invertebrates from the Niobrara River at Agate Fossil Beds National Monument, Nebraska using three methods and two sample handling protocols. We compared aquatic invertebrate assemblages collected with a Hester-Dendy multi-plate sampler, Hess sampler and a D-frame dipnet. We calculated six common bioassessment metrics from composite (combined) and replicate (separate) samples. Hess samples contained the highest taxonomic richness (capturing 77% of all taxa observed) and dipnet samples the least (47%). Hester-Dendy samples had the greatest proportion of Ephemeroptera, and Ephemeroptera, Plecoptera and Trichoptera (EPT). Dipnet samples had the lowest evenness values. In terms of sample handling, composite samples had inflated richness, diversity and evenness compared to replicate samples, but bioassessment metrics calculated from proportions or averages (i.e. Hilsenhoff’s Biotic Index and the proportion of EPT taxa) did not differ between them. The proportion of invertebrate groups from composite samples were not statistically different among sampling methods, but several groups differed between replicate samples collected by different methods. Ultimately, we recommend collecting replicate samples with a Hess sampler when the goal of the study is to detect ecosystem change, among locations or differences in variables of interest.

## Introduction

Aquatic invertebrates have been used to monitor ecosystem quality for over 150 years (Cairns and Pratt 1993), largely because they have several characteristics that make them ideal for the task. Aquatic invertebrates are relatively long lived (weeks to >100 years, Rosenberg and Resh 1993a) and unlike water samples that are collected periodically, invertebrates are permanent stream residents and therefore their presence or absence reflects long-term conditions at a site. For instance, water samples may miss discrete, short-lived discharges of pollution, but aquatic invertebrate communities will respond to such an event (Rosenberg and Resh 1993b). Furthermore, aquatic invertebrates are relatively sedentary, diverse and are inexpensive to collect and identify. Most importantly, lower ecosystem quality in a stream can increase mortality and decrease reproduction, survival and fitness of sensitive aquatic invertebrates (e.g., Ephemeroptera) whiles others are more tolerant to disturbances (e.g., Diptera; Johnson et al. 1993; Barbour et al. 1999). Changes in the diversity or assemblage structure of aquatic invertebrates can inform managers of stream ecosystem quality (Rosenberg and Resh 1993b).

Choosing a sampling method for aquatic invertebrate monitoring is difficult and depends on many variables. All approaches have advantages and disadvantages (e.g., cost to implement, time, bias towards specific taxa or life histories; e.g., Macanowicz et al. 2013, Tronstad and Hotaling, 2017). Therefore, identifying a method that is cost-effective, minimizes bias and fulfills management objectives is critical. Bioassessment studies use a variety of sampling methods, including kicknets, fixed-area samplers (e.g., Hess sampler), artificial substrates (e.g., Hester-Dendy samplers) and dipnets (Carter and Resh 2001). However, some sampling methods are not well-suited to all stream habitats. For example, artificial substrates (e.g., Hester-Dendy plates) are ideal for large, deep rivers that are otherwise difficult to sample (De Pauw et al. 1986). However, artificial substrates rely on colonization and therefore, do not represent natural assemblages or densities and can be biased towards certain insect orders (Letovsky et al. 2012). The type of information being collected also matters. For example, qualitative data may be sufficient if the study is estimating ecosystem health to meet federal standards, but more rigorous quantitative sampling is needed to assess change over time (e.g., Slavik et al. 2004). Qualitative samples only report proportional data, while fixed area samplers provide quantitative information on the density and biomass for each taxon in the assemblage.

Laboratory protocols can alter the taxa identified and the bioassessment metrics calculated. Previous studies (e.g., Vinson & Hawkins 1996) have investigated what type of subsampling method is best for bioassessment studies to minimize cost and produce reliable results. The two main types of subsampling – fixed area (e.g., 25% of sample) and fixed count (e.g., 300 individuals; e.g., King and Richardson 2002) – have been compared for many data types (e.g., Vinson & Hawkins 1996). However, the question of how replicate samples should be handled i.e., whether combined into composites or processed as replicates, remains largely unaddressed. Most bioassessment protocols (e.g., US EPA) direct users to composite samples in the field. That is, individual samples are combined into one large sample which is assumed to homogenize variance (Carey and Keough 2002); however, we are not aware of any studies investigating that assumption. Alternatively, replicate samples can be kept and analyzed separately with potential for added insight at relatively little additional cost. Replicate samples have rarely been integrated into bioassessment methods but a few exceptions occur. DiFranco (2014) recommends collecting three replicate samples in wetland habitats. Lazorchak et al (1998) and Hering et al. (2004) straddle a grey area between replicate and composite samples by directing users to pool microhabitat samples (e.g., pools and riffles) so that variance among habitats is estimated.

The National Park Service (NPS) has been monitoring aquatic invertebrates in the Niobrara River at Agate Fossil Beds National Monument since 1989 using Hester-Dendy samplers. However, due to the inherent complications of collecting samples using artificial substrates and an inability to make direct comparisons to other streams, a change in monitoring approach is under consideration. In this study, we used the opportunity to address an applied issue in stream biomonitoring and answer three questions: 1.) How does sampling method affect the invertebrate assemblage collected in the Niobrara River? 2.) How do the corresponding bioassessment metrics compare among sampling methods? And, 3.) to what degree do composite vs. replicate samples alter the assemblage and bioassessment metrics?

## Materials and methods

### Study area

The headwaters of the Niobrara River are located near Lusk, Wyoming and the river flows eastward into Nebraska and eventually into the Missouri River near Niobrara, Nebraska (Fig. 1). The Niobrara River Basin covers 32,600 km^2^ of which the majority is grassland in northern Nebraska (Galat et al. 2005). Over 95% of the land within the basin is used for agriculture. The Niobrara River flows through Agate Fossil Beds National Monument in western Nebraska about 23 km from the Wyoming border. Here, the Niobrara River is a low order stream flowing through grassland. Agate Fossil Beds National Monument includes ∼10.9 km^2^ in a valley bottom and ∼18 km of river flows through the park (Fig. 1). The river’s riparian vegetation is dominated by cattails (*Typha* sp.) and the invasive yellow flag iris (*Iris pseudacorus*) and its substrate is predominantly fine particles (e.g., sand, silt and clay). Currently, northern pike (*Esox lucius*), white suckers (*Catostomus commersonii*) and green sunfish (*Lepomis cyanellus*) inhabit the river within the park (Spurgeon et al. 2014); however, nine other fish species were collected at Agate Fossil Beds National Monument prior to 1990 (Spurgeon et al. 2014).

**Fig. 1.**
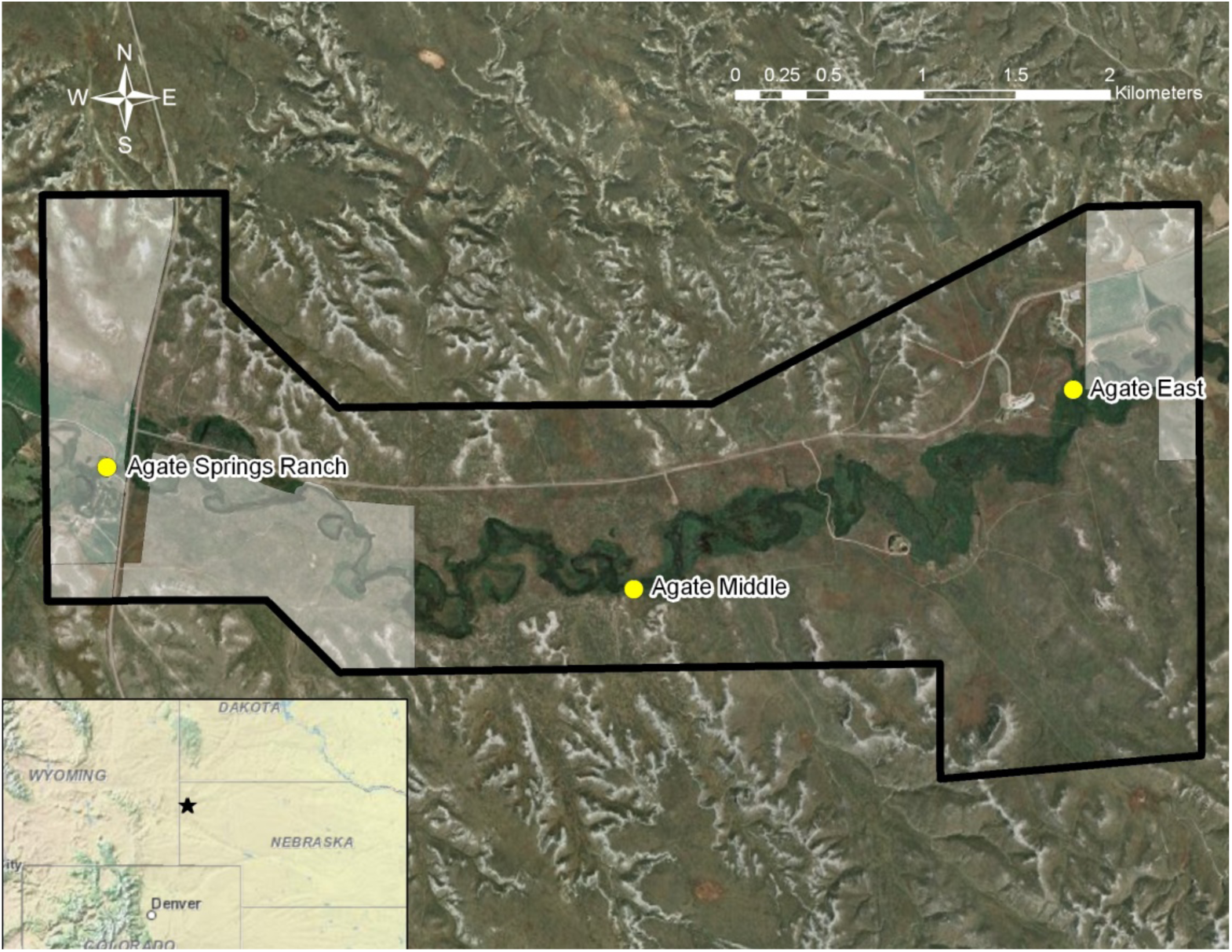
We sampled three sites along the Niobrara River at Agate Fossil Beds National Monument in Nebraska, USA. The black line is the Monument boundary and the transparent white areas are private land within the Monument. The inset shows the location of Agate Fossil Beds National Monument in Nebraska (star).

We sampled three long-term monitoring sites along the Niobrara River (Fig. 1; Tronstad & Hotaling, 2017) in 2016. We deployed Hester-Dendy samplers in mid-July and returned to collect them as well as Hess and dipnet samples in mid-August (see below). The most upstream site (Agate Springs Ranch) is located near the western park boundary. Agate Springs Ranch has an overstory of plains cottonwood (*Populus deltoides*) and cattails are more abundant than iris. The central site, Agate Middle, lacks an overstory and has gravel substrate with abundant iris and cattails surrounding the river. Finally, Agate East is located before the Niobrara River flows out of the park and is the deepest site with riparian vegetation dominated by iris and a few willows (*Salix* spp.).

### General measurements

To assess general environmental characteristics of our study sites, we measured a number of standard variables (e.g., temperature), as well as water quality and clarity, sediment composition, water depth and discharge. We measured dissolved oxygen (percent saturation and mg/L), pH, water temperature, specific conductivity and oxidation-reduction potential using a Yellow Springs Instruments (YSI) Professional Plus. The YSI was calibrated on-site before use. We measured water clarity by estimating the depth at which a Secchi disk disappeared from sight. The dominant substrate was recorded in the main channel of all sites and where each Hess sample was taken using soil texture tests (Thien 1979). Clay was defined as fine particles forming a ribbon after removing water, whereas silt did not form a ribbon. Sand was characterized by particles 0.06-2 mm in diameter, gravel was 2-64 mm in diameter, cobble was 64-256 mm in diameter, boulders were 25-400 cm in diameter, bedrock was >4 m in diameter and hardpan/shale was identified by firm, consolidated fine substrate. We recorded the location of each site using a global positioning system (GPS; Garmin eTrex Vista HCx). Finally, we estimated stream discharge (*Q*; m^3^/s) by measuring water depth (*d*; m) and velocity (*v*; m/s) using a Marsh-McBirney Flo-Mate 2000 at 0.3 m intervals across the stream’s width (*w*; m) and summing each interval using Equation 1:

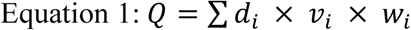

### Hester-Dendy sample collection

We deployed seven Hester-Dendy samplers (76 mm x 76 mm, 9 plates, Wildlife Supply Company) at each site. For each sampler, we strung a rope across the stream between two fixed posts with evenly spaced loops to separate the Hester-Dendy multiplate samplers. The Hester-Dendy samplers were suspended in the water column at least 15 cm above the substrate. Debris dams were cleared weekly and we retrieved the samplers after 30 days of colonization by approaching the site from downstream, placing a dipnet (150 µm mesh) under it and cutting the rope. Hester-Dendy samplers were immediately placed in a container with ∼80% ethanol and any organisms in the dipnet were removed and placed in the same container. In the laboratory, we dismantled and scrubbed the Hester-Dendy samplers to remove invertebrates that colonized the plates, then we rinsed the samplers through a 212 µm sieve and preserved all specimens in ∼80% ethanol. The middle five Hester-Dendy samples were used for analysis except when one of the samplers were compromised (e.g., touching the bottom).

### Hess sample collection

We collected five Hess samples (500 µm mesh, 860 cm^2^ sampling area, Wildlife Supply Company) at each site. Samples were taken along the shallower margins of the stream where emergent vegetation is abundant. We placed the Hess sampler over vegetation to collect invertebrates living on it and in the surrounding benthic sediment. The vegetation and sediment were vigorously agitated and invertebrates were captured in the net. Samples were preserved in 80% ethanol and returned to the laboratory for analysis.

### Dipnet sample collection

We collected dipnet samples along a reach that was 40x the wetted stream width following standard methods for sampling aquatic invertebrates in wadeable streams (US EPA 2013). We measured the wetted width at five representative points along the stream and averaged values to the nearest meter. The average width of the Niobrara River was less than 4 m, so we used a minimum reach length of 150 m. We sampled invertebrates along 11 evenly-spaced transects that were 15 m apart using a D-frame net (243 µm mesh, 30.5 × 25.4 cm opening, Wildlife Supply Company). At each transect, we sampled the right, left and center of the stream systematically. Multiple habitats were sampled including benthic substrate, woody debris, macrophytes and leaf packs. All samples were composited and preserve in the field with 95% ethanol.

For dipnet sampling, we classified streams into riffle/run or pool/glide habitat and adjusted our methods for each. We defined a habitat as riffle/run if the current fully extend the net or a pool/glide if the net did not fully extend. For riffle/run habitats, we placed the net on the bottom of the stream with the opening facing upstream. We visually defined a sampling area as one net width wide and long upstream of the opening (∼30 × 25 cm). We first removed any large organisms (e.g., snails, mussels) from the sampling area and placed them into the net. Next, we scrubbed all rocks that were golf ball sized (∼4 cm) or larger to dislodge organisms, wash them into the net and placed the scrubbed rocks outside of the sampling area. Finally, we held the net below the sampling area and disturbed the remaining finer substrate for 30 seconds while the drift washed into the net. Pool/glide habitats were sampled the same as riffle/run except the net was repeatedly pulled through the disturbed water just above the substrate to capture organisms and continuously moved throughout sampling to ensure no organisms escaped the net.

After we sampled a transect, we transferred the sample to a sieve bucket (500 µm mesh). We removed as much gravel as possible and inspected the net for any residual organisms. We inspected each large object (e.g., rocks or sticks), removed organisms that were attached to them and discarded the object. For each sampled area, we recorded the dominant substrate size (e.g., fine/sand, gravel, coarse, other) and the habitat type (riffle/run or pool/glide).

### Sample processing – Hester-Dendy and Hess

Invertebrates collected with Hester-Dendy and Hess samplers were sorted from debris in white trays and identified under a dissecting microscope. We rinsed all samples through a 2 mm sieve followed by 212 µm (Hester-Dendy) or 500 µm (Hess) sieves to separate larger and smaller invertebrates. All large invertebrates (> 2 mm) were identified. If invertebrates were visually numerous in the smaller sieve, we subsampled the contents using the record player method (Waters 1969). Invertebrates were identified according to Merritt et al. (2008) for insects, and Thorp and Covich (2010) and Smith (2001) for non-insect invertebrates. Invertebrate tolerance values were assigned to each taxon from Barbour et al. (1999).

### Sample processing - Dipnet

We processed dipnet samples following the official EPA protocol (US EPA 2013). We elutriated all dipnet samples to remove inorganic substrate with a 500 µm mesh sieve. In the laboratory, we spread the sample evenly over a 30 × 36 cm sorting tray that was divided into 30 numbered grids (6 cm^2^ each). Using a random number generator in R (R Development Core Team 2013), we selected six of the 30 grids, removed the invertebrates and counted them. If the first six grids did not contain a minimum of 500 individuals, we randomly selected additional grids until the minimum threshold was reached. We removed and identified large or rare invertebrates defined as longer than 1.2 cm (Vinson and Hawkins 1996). All invertebrates were identified to the lowest taxonomic level possible, typically genus, and we normalized our abundance estimates for each site based upon the number of grids that were counted.

### Statistical analyses

We used R (R Development Core Team 2013) and the packages *plyr* (Wickham 2011), *Matrix* (Bates and Maechler 2013), and *vegan* (Oksanen et al. 2013) to calculate invertebrate abundances, proportions, bioassessment metrics and perform statistical tests. To estimate ecosystem quality, we calculated six common bioassessment metrics: Hilsenhoff’s Biotic Index (HBI), Ephemeroptera, Plecoptera and Trichoptera (EPT) richness, proportion of EPT taxa (number of EPT taxa divided by the total number of taxa collected), taxonomic diversity (Shannon’s index), taxonomic richness and taxonomic evenness.

We compared invertebrate proportions and bioassessment metrics among sites and sampling methods with ANOVAs. If sites or methods were significantly different, we used Tukey’s honest significant difference (HSD) to verify which sites or methods differed from one another with pair-wise comparisons. To compare invertebrate assemblages recovered with Hester-Dendy and Hess samples to dipnet samples, we electronically composited replicates at each site. However, to explore how compositing samples affects bioassessment metrics, we also calculated bioassessment metrics separately for each Hester-Dendy and Hess replicate at each site.

We evaluated differences in the aquatic invertebrate assemblage across sites and sampling method with non-metric multidimensional scaling (NMDS) implemented in the R package *vegan* (Oksanen et al. 2013). NMDS provides an ordination-based approach to rank distances between objects and has been shown to perform well with non-normally distributed data (Legendre and Legendre 1998). To prepare our data for NMDS analysis, we removed rare taxa (as defined as any taxon that was unique to a single site+method combination). Next, we calculated the mean and standard deviation (SD) for each taxon and removed two species which were present at more than two deviations above the mean. Finally, we removed any taxon present at less than 0.1% of the overall abundance (after the first two filtering steps were completed). NMDS analyses were performed using Bray-Curtis distances on composite samples with default settings. To test whether the assemblages recovered were different depending on sampling method or site, we performed an analysis of similarities (ANOSIM) with default settings (including 999 permutations). Next, we investigated differences in multivariate dispersion for each method by calculating the mean distance of each sample to the group’s centroid in multivariate space with the function *betadisper*. We assessed pair-wise differences in dispersion with a Tukey’s HSD. To better visualize taxonomic differences in invertebrate assemblages collected with each sampling method, we constructed a ternary plot using the R package *ggtern* (Hamilton 2015). For ternary plot construction, we only removed rare taxa (as described above) before averaging the abundances of each taxon in composite samples across sites for each method.

## Results

### Environmental variation

Sites were environmentally similar to one another with little variation between our July and August sampling dates (Table 1). Water temperatures ranged from ∼21-24°C. Dissolved oxygen concentrations were near saturation. Specific conductivity was approximately 350 µS/cm and pH was consistently highest at Agate Springs Ranch. Oxidation-reduction potential was highest at Agate Springs Ranch (169-197 mV) and we measured reducing conditions (< 200 mV) at all sites. Discharge was higher in August and Agate East had the lowest flow. Agate East was the deepest site (1.2-1.5 m). Agate Springs Ranch was the narrowest (3-3.8 m) and shallowest (0.5-0.7 m; Table 1) site. The substrate at all sites was dominated by fine sediment (i.e., clay, sand and silt) and gravel.

**Table 1.** Water quality and site characteristics measured when Hester-Dendy samplers were deployed (July) and when Hester-Dendy, Hess and dipnet samples were collected (August). A “B” after the Secchi disk depth indicated that the bottom of the stream was visible and the number is the maximum depth at the site. Stream width was measured with emergent vegetation excluded. Abbreviations and units include: T_WATER_ = water temperature, T_AIR_ = air temperature, DO = dissolved oxygen, SPC = specific conductivity, and ORP = oxidation-reduction potential.

| Parameter | Ranch | Middle | East | Ranch | Middle | East |
| --- | --- | --- | --- | --- | --- | --- |
| Date | 18 July | 18 July | 19 July | 19 Aug | 17 Aug | 17 Aug |
| Time | 13:50 | 18:00 | 11:15 | 13:15 | 15:30 | 17:15 |
| T<sub>WATER</sub> (°C) | 23.8 | 21.1 | 21.6 | 21.7 | 21.1 | 22.9 |
| T<sub>AIR</sub> (°C) | 30 | 28 | 30 | 34 | 36 | 28 |
| DO (% sat.) | NA | NA | NA | 107.0 | 98.0 | 107.0 |
| DO (mg/L) | NA | NA | NA | 8.0 | 7.3 | 7.9 |
| SPC (μS/cm) | 357.2 | 352.4 | 364.9 | 347.2 | 354.4 | 358.6 |
| pH | 8.5 | 8.1 | 7.9 | 8.5 | 8.0 | 8.2 |
| ORP (mV) | 168.7 | 45.2 | 32.5 | 196.6 | 72.6 | 81.1 |
| Secchi depth (cm) | 47 (B) | 82 (B) | 67 (B) | 58.5 (B) | 73 (B) | 149.0 |
| Max. depth (m) | 1.6 | 2.7 | 4.0 | 2.2 | 2.4 | 4.9 |
| Width (m) | 12.4 | 14.0 | 12.7 | 9.7 | 13.5 | 16.4 |
| Discharge (m<sup>3</sup>/s) | 0.18 | 0.21 | 0.13 | 0.22 | 0.27 | 0.17 |
| Substrate | Sand | Gravel | Silt | Sand | Gravel | Silt/sand |

### Community composition

We identified 73 invertebrate taxa representing six phyla (Annelida, Arthropoda, Mollusca, Nematoda, Nematomorpha and Platyhelminthes) in the Niobrara River when all samplers were combined (SM A-C). Hester-Dendy samples contained nine taxa not found in Hess samples, 18 taxa not collected with the dipnet and 8 taxa unique to Hester-Dendy samples. Hess samples contained 30 taxa not collected with Hester-Dendy samplers, 31 taxa not collected with the dipnet and 21 taxa unique to Hess samples. Dipnet samples included 16 taxa not collected with Hester-Dendy samplers, 10 taxa not present in Hess samples and 8 taxa unique to dipnet samples.

When composited, proportions of insects (Fig. 2a; *F* = 0.3, *df* = 1, *p* = 0.75) and non-insects (Fig. 2b; *F* = 0.3, *df* = 1, *p* = 0.75) did not differ among sampling methods. Proportions of Annelida, Crustacea, Coleoptera, Diptera, Ephemeroptera, Hemiptera, Mollusca, Odonata and Trichoptera also did not differ when composited (*p* ≥ 0.25; Fig. 2). Conversely, when treated as replicates, the proportion of insects (Fig. 2a; *F* = 4.8, *df* = 1, *p* = 0.04), non-insects (Fig. 2b; *F* = 4.8, *df* = 1, *p* = 0.04), Annelida (Fig. 2c; *F* = 11.8, *df* = 1, *p* = 0.002), Ephemeroptera (Fig. 2d; *F* = 4.6, *df* = 1, *p* = 0.04), Odonata (Fig. 2e; *F* = 4.6, *df* = 1, *p* = 0.04) and Trichoptera (Fig. 2f; *F* = 6.9, *df* = 1, *p* = 0.01) differed between Hester-Dendy and Hess samples. The proportion of Mollusca (*F* = 3.7, *df* = 1, *p* = 0.065), Crustacea (*F* = 0.43, *df* = 1, *p* = 0.52), Coleoptera (*F* = 0.2, *df* = 1, *p* = 0.65), Diptera (*F* = 0.79, *df* = 1, *p* = 0.38) and Hemiptera (*F* = 2.5, *df* = 1, *p* = 0.13) did not differ between replicate Hester-Dendy and Hess samples.

**Fig. 2.**
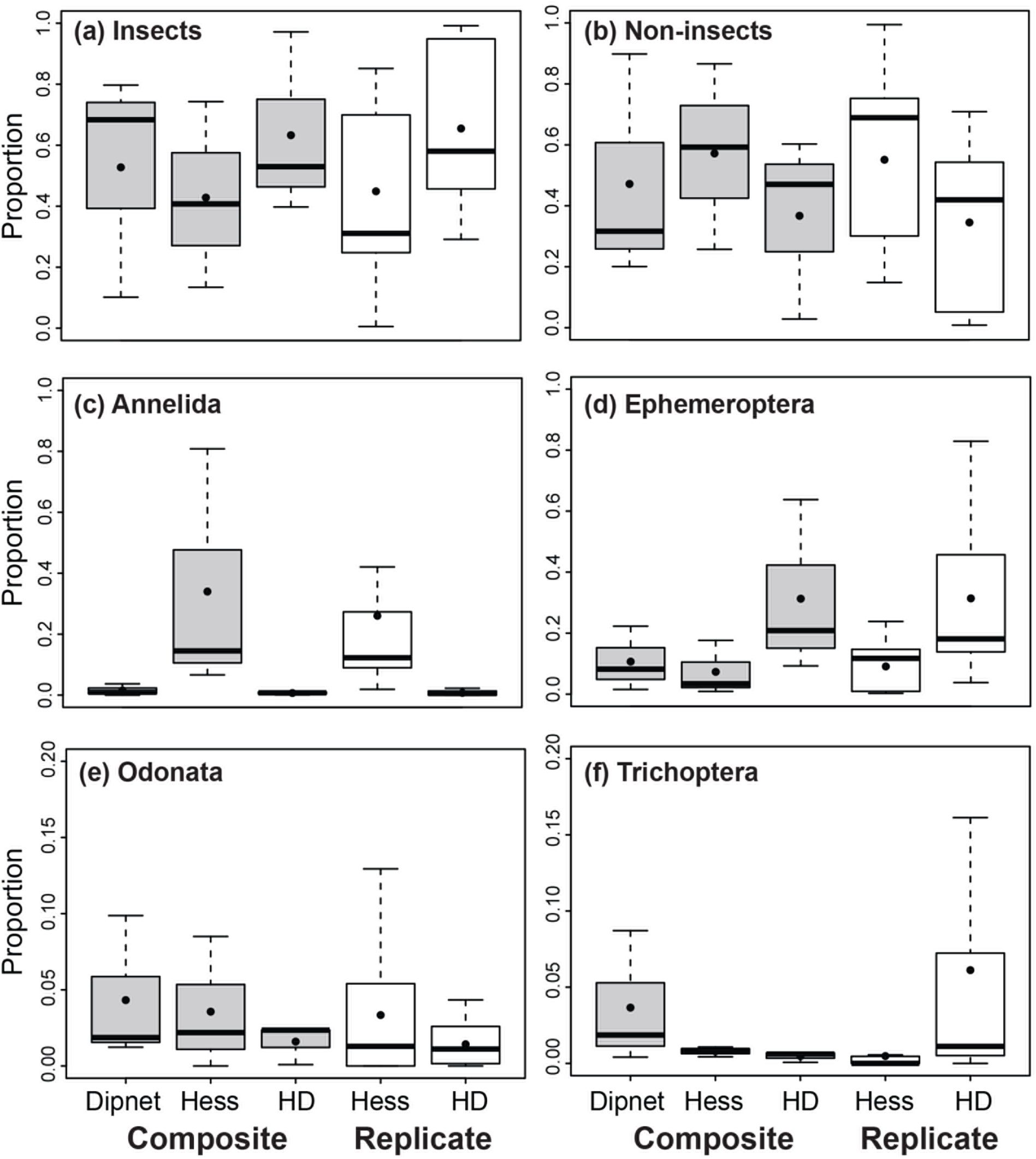
Proportions of insects (a), non-insect invertebrates (b), Annelida (c), Ephemeroptera (d), Odonata (e) and Trichoptera (f) in dipnet, Hess and Hester-Dendy (HD) samples that were composited (grey boxes) or kept separate as replicates (white boxes; HD and Hess only) collected from the Niobrara River, Nebraska, USA. Black circles are mean values, bold lines are median values, lower and upper limits are the 25^th^ and 75^th^ percentiles and whiskers indicate the lower and upper limits of the data.

Additionally, NMDS analyses indicated that the sampling methods collected different aquatic invertebrate assemblages (*p*, ANOSIM = 0.008; Fig. 3a), but that overall, assemblages did not differ among sites (*p*, ANOSIM = 0.408; Fig. 3b). While different sampling methods yielded distinct assemblages, the amount of multivariate space occupied by each method did not differ (*p*, Tukey’s HSD ≥ 0.94). Visualization of the assemblage recovered by each method via ternary plot highlighted the strong bias towards Hess and Hester-Dendy sampling in terms of unique taxa (Fig. 4). After filtering rare taxa as described above, only one taxon, *Ceratopogon*, a genus of Ceratopogonidae, was observed in dipnet samples yet was largely absent elsewhere. Both Hess (13 taxa) and Hester-Dendy (7 taxa) sampling recovered a number of taxa that were either rare or completely absent in the results of the other methods. However, some taxa were relatively equally represented across all three methods including *Anax*, Collembola, *Hyallela* and Lymnaeidae (Fig. 4).

**Fig. 3.**
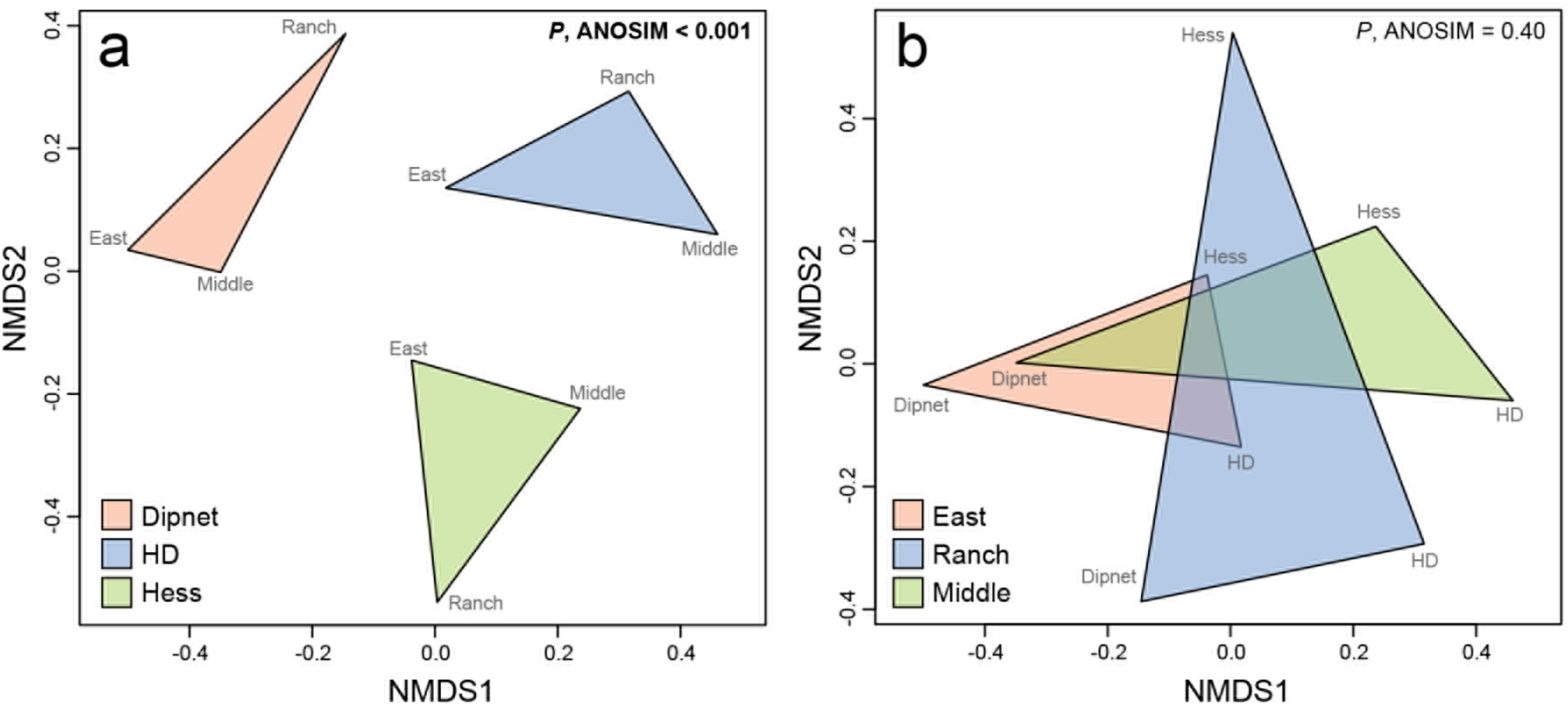
Comparisons of invertebrate assemblages recovered by (a) sampling method and (b) site with non-metric multidimensional scaling (NMDS). Collected assemblages differed with sampling method but not site. HD = Hester-Dendy.

**Fig. 4.**
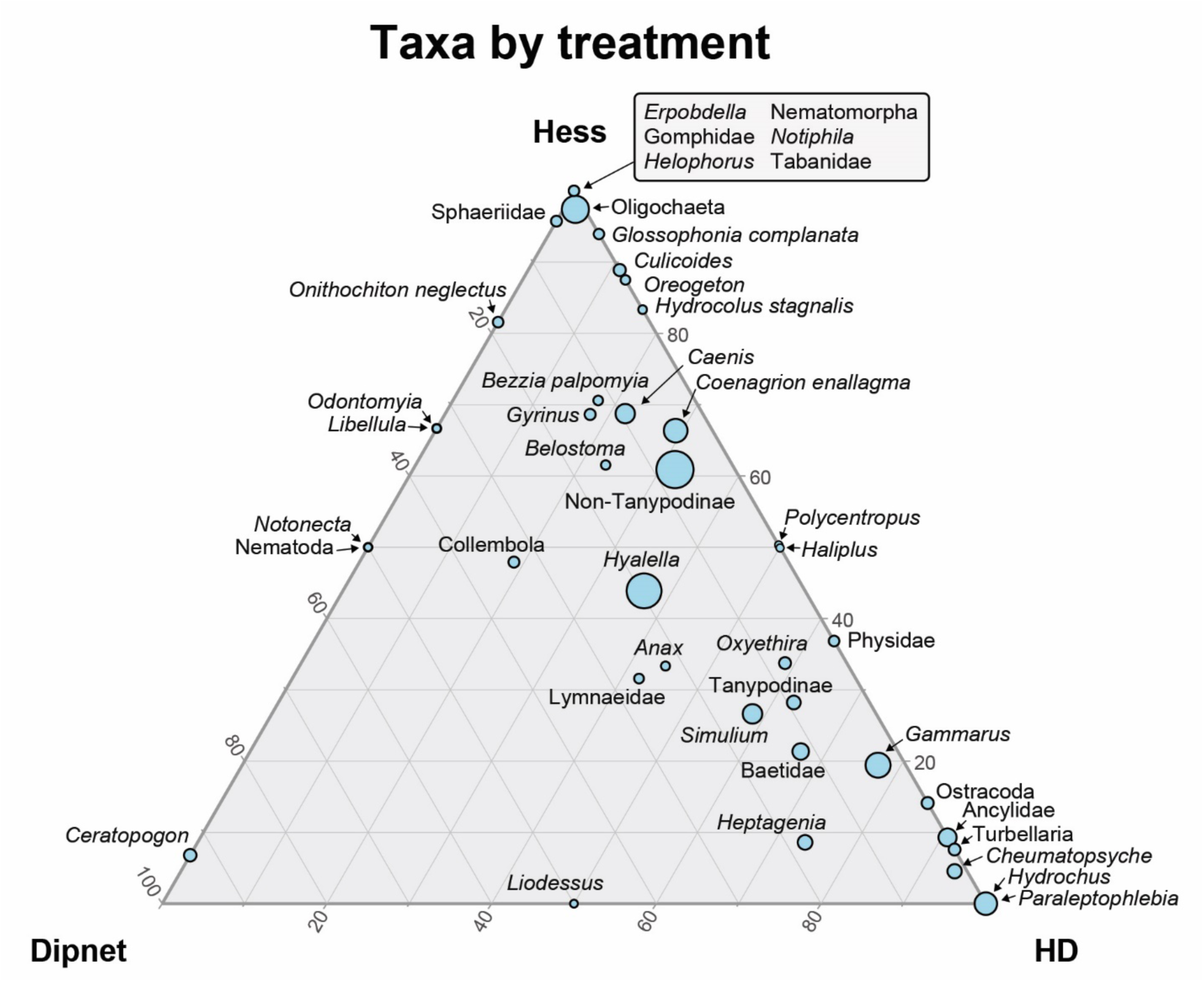
Distribution of taxa recovered by Hess, Hester-Dendy and dipnet sampling in the Niobrara River. The position of a given point indicates the percentage of the associated taxon with each sampling method. Circle size indicates the relative abundance of each taxon overall.

### Bioassessment metrics

When calculated from composite samples, bioassessment metrics differed among sampling methods, but most comparisons were not significant without incorporating replicates. Taxonomic richness (Fig. 5a; *F* = 2.6, d*f* = 2, *p* = 0.19), diversity (Fig. 5b; *F* = 4.4, d*f* = 2, *p* = 0.10), evenness (Fig. 5c; *F* = 5.4, d*f* = 2, *p* = 0.07) and EPT richness (Fig. 5d; *F* = 3.3, d*f* = 2, *p* = 0.14) did not differ among sampling methods. The proportion of EPT taxa (Fig. 5e; *F* = 63, d*f* = 2, *p* = 0.0009) were highest in Hester-Dendy samples and lowest in Hess samples (Tukey’s HSD, *p* < 0.05). HBI values (Fig. 5f; *F* = 28, d*f* = 2, *p* = 0.005) were lower in Hester-Dendy samples (Tukey’s HSD, *p* < 0.02).

**Fig. 5.**
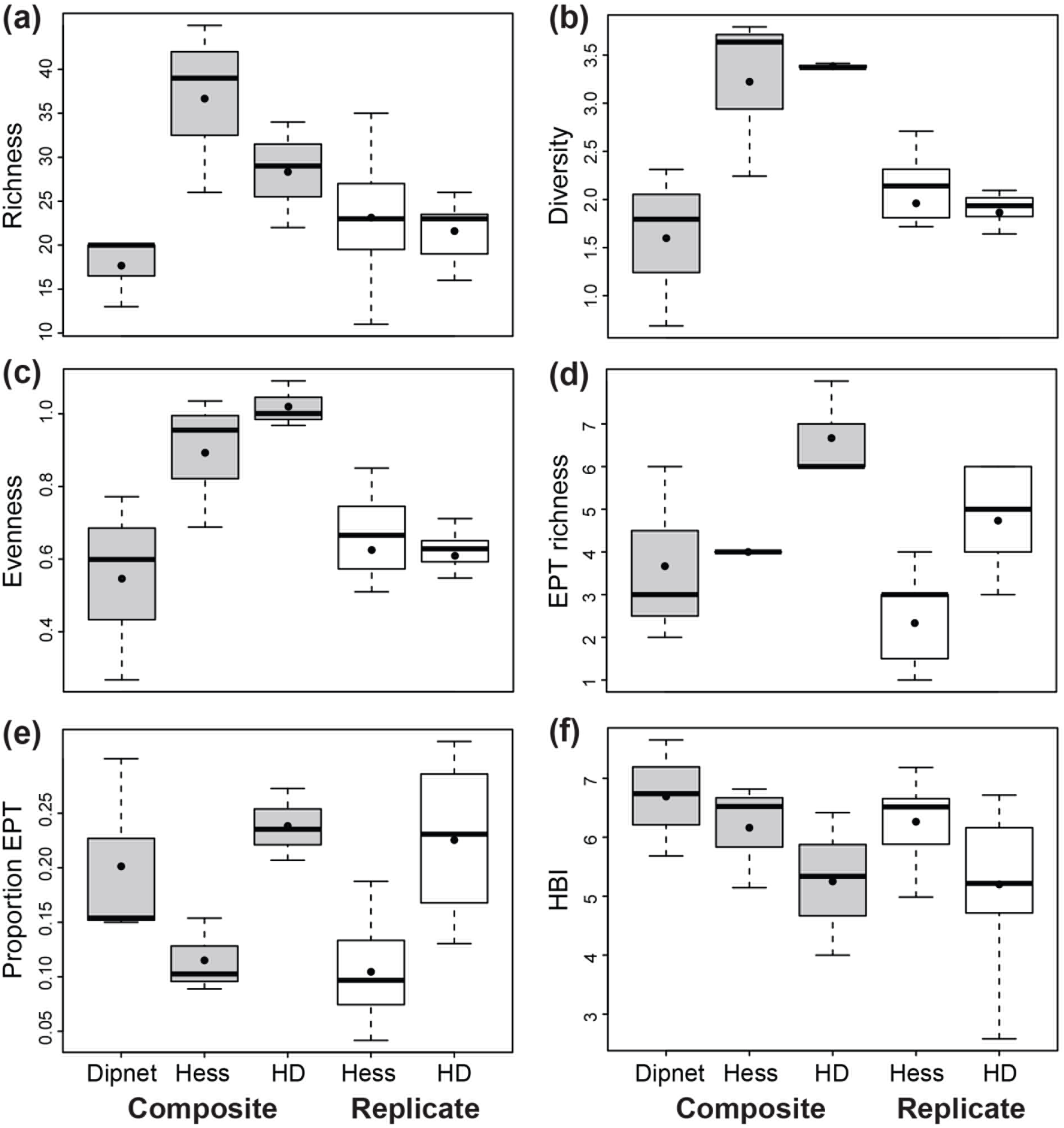
(a) Richness, (b) diversity, (c) evenness, (d) Ephemeroptera, Plecoptera and Trichoptera (EPT) richness, (e) proportion of EPT taxa and (f) Hilsenhoff’s biotic index (HBI) calculated from dipnet, Hester-Dendy (HD) and Hess samples for this study. Metrics calculated from composited samples are in grey and those calculated from five replicate samples are in the white boxes. For all metrics, except HBI, higher values indicate better ecosystem quality. Black circles represent mean values and bold lines are median values, lower and upper edges of the box are the 25^th^ and 75^th^ percentiles and whiskers indicate the lower and upper limits of the data.

Most bioassessment metrics calculated from electronically composited samples were higher than those estimated from replicate samples. When composited, 40% and 80% more taxa were observed in Hester-Dendy and Hess samples, respectively, versus replicate samples (Table 2). Similarly, EPT richness was 43% and 83% higher in composited Hester-Dendy and Hess samples, respectively, versus replicates. Taxonomic diversity was also 82% higher in composited Hester-Dendy samples and 63% higher in composited Hess samples. Finally, composited Hester-Dendy and Hess samples had 58% and 54% higher evenness values, respectively. Conversely, the proportion of EPT taxa and HBI values did not differ between composite and replicate samples.

**Table 2.**
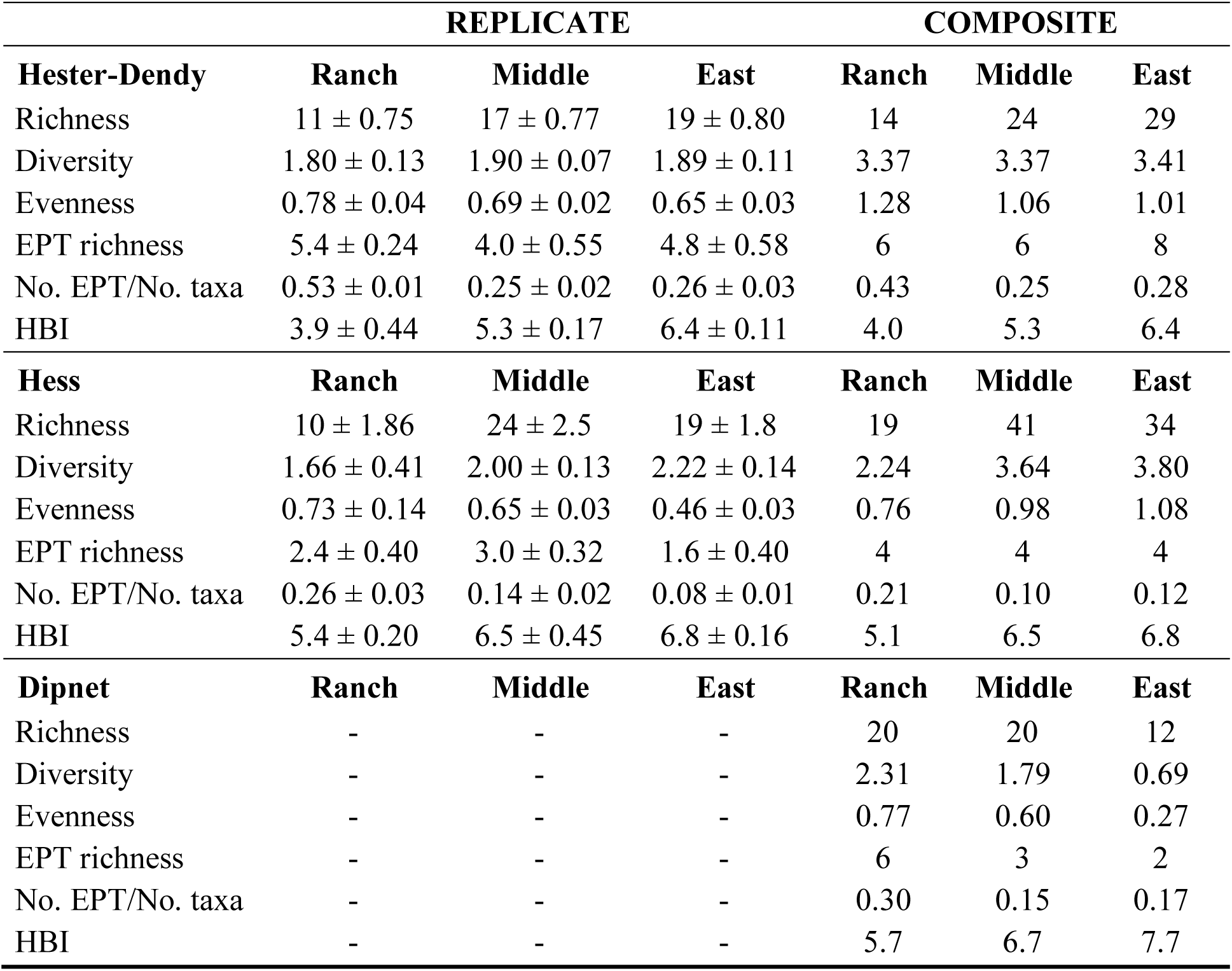
Invertebrate bioassessment metrics calculated from Hester-Dendy, Hess and dipnet samples collected in the Niobrara River. Metrics for Hester-Dendy and Hess samples were calculated from replicate samples (i.e., mean metrics ± standard error) and composited samples (all replicate samples combined for each site and sampler). Dipnet samples were composited in the field and therefore no replicate samples are available for comparison.

|  | REPLICATE |  |  | COMPOSITE |  |  |
| --- | --- | --- | --- | --- | --- | --- |
| **Hester-Dendy** | **Ranch** | **Middle** | **East** | **Ranch** | **Middle** | **East** |
| Richness | 11 ± 0.75 | 17 ± 0.77 | 19 ± 0.80 | 14 | 24 | 29 |
| Diversity | 1.80 ± 0.13 | 1.90 ± 0.07 | 1.89 ± 0.11 | 3.37 | 3.37 | 3.41 |
| Evenness | 0.78 ± 0.04 | 0.69 ± 0.02 | 0.65 ± 0.03 | 1.28 | 1.06 | 1.01 |
| EPT richness | 5.4 ± 0.24 | 4.0 ± 0.55 | 4.8 ± 0.58 | 6 | 6 | 8 |
| No. EPT/No. taxa | 0.53 ± 0.01 | 0.25 ± 0.02 | 0.26 ± 0.03 | 0.43 | 0.25 | 0.28 |
| HBI | 3.9 ± 0.44 | 5.3 ± 0.17 | 6.4 ± 0.11 | 4.0 | 5.3 | 6.4 |
| **Hess** | **Ranch** | **Middle** | **East** | **Ranch** | **Middle** | **East** |
| Richness | 10 ± 1.86 | 24 ± 2.5 | 19 ± 1.8 | 19 | 41 | 34 |
| Diversity | 1.66 ± 0.41 | 2.00 ± 0.13 | 2.22 ± 0.14 | 2.24 | 3.64 | 3.80 |
| Evenness | 0.73 ± 0.14 | 0.65 ± 0.03 | 0.46 ± 0.03 | 0.76 | 0.98 | 1.08 |
| EPT richness | 2.4 ± 0.40 | 3.0 ± 0.32 | 1.6 ± 0.40 | 4 | 4 | 4 |
| No. EPT/No. taxa | 0.26 ± 0.03 | 0.14 ± 0.02 | 0.08 ± 0.01 | 0.21 | 0.10 | 0.12 |
| HBI | 5.4 ± 0.20 | 6.5 ± 0.45 | 6.8 ± 0.16 | 5.1 | 6.5 | 6.8 |
| **Dipnet** | **Ranch** | **Middle** | **East** | **Ranch** | **Middle** | **East** |
| Richness | - | - | - | 20 | 20 | 12 |
| Diversity | - | - | - | 2.31 | 1.79 | 0.69 |
| Evenness | - | - | - | 0.77 | 0.60 | 0.27 |
| EPT richness | - | - | - | 6 | 3 | 2 |
| No. EPT/No. taxa | - | - | - | 0.30 | 0.15 | 0.17 |
| HBI | - | - | - | 5.7 | 6.7 | 7.7 |

### Hester-Dendy sampling

Across all methods and sites, Hester-Dendy samples contained 52% of the total invertebrate community we observed. Insecta and Crustacea (90% of individuals) were the most abundant taxa in Hester-Dendy samples. Of the insects, Diptera and Ephemeroptera were the most abundant followed by Trichoptera and Odonata (SM 1). Hester-Dendy samples from Agate Middle (909 ind/sample) contained more invertebrates than both Agate Springs Ranch (217 ind/sample) and Agate East (279 ind/sample; *F* = 7.1, *df* = 2, *p* = 0.009; Tukey HSD, *p* < 0.025; calculated with replicate samples). Taxonomic richness was lowest at Agate Springs Ranch (Table 2; *F* = 28.7, *df* = 2, *p* < 0.001). Taxonomic diversity (*F* = 0.35, *df* = 2, *p* = 0.71), taxonomic evenness (*F=* 0.25, *df* = 2, *p* = 0.78), EPT richness (Table 2; *F* = 2.1, *df* = 2, *p* = 0.16) and the proportion of EPT taxa did not differ among sites (Table 2; *F* = 1.8, *df* = 2, *p* = 0.2). The average tolerance value for an invertebrate collected with Hester-Dendy sampling was lowest at Agate Springs Ranch (HBI; Table 2; *F* = 18.9, *df* = 2, *p* < 0.001; Tukey HSD, *p* ≤ 0.05).

### Hess sampling

We collected 77% of all observed taxa with Hess sampling. Overall, Insecta, Crustacea and Annelida (98% of individuals) were the most numerous groups in Hess samples. Of the insects, Diptera were most abundant followed by Ephemeroptera, Odonata and Trichoptera (SM 2). Hess samples from Agate Middle (926 ind/sample) had higher abundances of invertebrates compared to both Agate East (465 ind/sample) and Agate Springs Ranch (282 ind/sample; *F* = 8.7, *df* = 2, *p* = 0.005; Tukey HSD, *p* ≤ 0.035; calculated from replicate samples). Taxonomic richness was lowest at Agate Springs Ranch (Table 2; *F* = 11.7, *df* = 2, *p* = 0.001; Tukey’s HSD, *p* < 0.02), but taxonomic diversity did not differ among sites (Table 2; *F* = 5.3, *df* = 2, *p* = 0.02). Taxonomic evenness was highest at Agate Springs Ranch (Table 2; F = 14.6, df = 2, p < 0.001; Tukey HSD, p ≤ 0.01). Agate Springs Ranch also had a higher proportion of EPT taxa than both other sites (Table 2; F = 3.8, df = 2, p = 0.05). Additionally, invertebrates at Agate Springs Ranch had the lowest mean tolerance value (HBI; Table 2; F = 24, df = 2, p < 0.0001; Tukey’s HSD, *p* < 0.001).

### Dipnet sampling

Of all the invertebrate taxa observed in this study, 47% were found in dipnet samples. Overall, Insecta and Crustacea (99% of individuals) were the most numerous invertebrates. Within insects, Diptera were the most abundant order followed by Ephemeroptera, Odonata and Coleoptera (SM 3). We collected the most individuals from Agate Middle (∼2685 ind/sample) and fewer individuals from Agate East (∼1260 ind/sample) and Agate Springs Ranch (∼400 ind/sample). Taxonomic richness and diversity were lowest at Agate East (Table 2). Taxonomic evenness was highest at Agate East (Table 2). Agate Springs Ranch had the highest number of EPT as well as the highest EPT proportion (Table 2). As a result, invertebrates at Agate Springs Ranch had the lowest mean tolerance value (HBI). No statistical comparisons among sites are reported due to the lack of replicates for the dipnet sampling.

## Discussion

Both sampling method and processing (whether replicate or composite) alters the invertebrate assemblage collected and bioassessment metrics calculated. Hess samples yielded more unique taxa and the most complete picture of the stream invertebrate assemblage. Hester-Dendy samples were biased toward EPT taxa and dipnet sampling emphasized the most common taxa and thus had the lowest evenness values. Compositing samples yields elevated taxonomic richness, diversity and evenness compared to the same metrics calculated from individual replicates; however, metrics based on proportions or averaging (e.g., HBI) did not differ. Our results add another line of evidence that different sampling methods collect different portions of the invertebrate community and care must be taken when choosing an approach. For example, many studies have compared the aquatic invertebrates captured using different samplers in a variety of habitats, such as streams, wetlands, vegetation and sink holes (e.g., Macanowics et al. 2013; Turner and Trexler 1997; Buss and Borges 2008); however, we are unaware of any studies comparing Hess, Hester-Dendy and dipnet sampling directly. While managers should be aware of the potential bias of different methods, some approaches may be more useful than others under certain conditions. For example, funnel traps, dipnets and stovepipe corers captured the most taxa in emergent vegetation of the Florida Everglades while Hester-Dendy sampling collected fewer taxa (Turner and Trexler 1997). Similar to the Niobrara River, quantitative Surber samplers (an analog of Hess sampling) collected 95-98% of taxa in two Australian rivers where qualitative kicknet samples only captured 63-66% of the community (Gillies et al. 2009).

Bioassessment metrics are also influenced by sampling method (e.g., Bouchard et al. 2014), sorting technique (e.g., Nichols and Norris 2006), subsampling method (e.g., Nichols and Norris 2006; King and Richardson 2002), mesh size (e.g., Battle et al. 2007) and the taxonomic level specimens are identified to (e.g., King and Richardson 2002; Jones 2008). Despite the fact that compositing samples is common in stream bioassessment (e.g., US EPA 2013, RIVPACS), few studies have investigated how compositing samples may alter metrics. We show that compositing alters bioassessment metrics (e.g., taxonomic richness, diversity and evenness) and therefore, metrics calculated from composite samples should not be compared to those calculated from replicate samples. Indeed, only metrics calculated from proportions or averages should be compared between composite and replicate samples.

Composite samples are typically used as a cost-efficient method to assess conditions in aquatic ecosystems when estimating variance is not critical (Downes 2010). Most bioassessment protocols (e.g., RIVPACS and US EPA) recommend compositing samples to calculate a single estimate of metrics per site. Collecting a large composite sample is presumed to homogenize the variance, and therefore produce a single, reliable value (Carey & Keough 2002; B. Marshall, personal communication). One study discovered that metrics calculated using composite samples varied by 30% within a site (B. Marshall, personal communication). Vlek et al. (2006) compared the ecological quality class (a measure of stream ecosystem health) from bioassessment metrics calculated with replicate and composite samples, and found that 8% were in different classes when five replicate samples were collected. In our study, composite samples from all methods produced a different result for each site using Hilsenhoff’s Biotic Index (Hilsenhoff 1987). Composited Hester-Dendy samples had the highest ratings (fair to very good) and dipnet samples the lowest (poor to fair). Bradley and Ormerod (2002) reported that rare taxa were the largest source of error when sampling streams with kicknets. Another source of error likely lies in subsampling of large composite samples which may introduce variance compared to replicate samples. Regardless of the subsampling method (i.e., fixed area or fixed counts), fewer individuals are removed and analyzed in composite samples versus replicate samples. Ultimately, more individuals analyzed will always yield more accurate estimates of conditions, but increasing the number of individuals also requires more resources. More studies designed to estimate differences between composite and replicate samples and their associated bioassessment metrics are needed to understand the consequences of sampling designs and when it’s appropriate to use them.

Unlike composite samples, replicates enable managers to calculate variance which provides additional power to estimate differences among variables and/or sites of interest while simultaneously improving bioassessment accuracy (Quinn and Keough 2002). A key to effective use of replicate samples lies in identifying the variables for which knowledge of the variance is valuable, and collecting replicates for them, while also identifying when to composite samples for other variables to save resources (Downes 2010). Replicate samples are recommended for monitoring data where statistical power is needed to detect changes over time (e.g., Slavik et al. 2004). Replicates are also necessary when the goal of a study is to detect differences among variables (e.g., sites, substrate), because replicates provide vital statistical power. For example, when replicates were composited in our study, we did not detect statistically significant differences in the proportion of invertebrate groups or the calculated metrics (e.g., taxonomic richness); however, when replicates for Hester-Dendy and Hess samples were compared, many groups yielded statistically different results. For best practices in stream biomonitoring, we recommend collecting replicate samples that are analyzed separately and electronically composited later if the need arises. While an argument could be made that collecting one composited sample in the field reduces the number of samples to manage in transit and process, in our experience, replicate samples are easier to process in the laboratory as they reduce the amount of material per sample, especially in areas with a lot of organic matter.

We also showed that different sampling methods yield very different perspectives on the aquatic invertebrate community being studied. Previous studies have reported that Hester-Dendy sampling tends to select for EPT taxa (Canton and Chadwick 1983; Letovsky et al. 2012). Because EPT richness is a common metric in biomonitoring, Hester-Dendy samples can bias bioassessment metrics towards lower values, indicating better ecosystem health. Our results support this as Hester-Dendy samples in the Niobrara River had the largest proportion of Ephemeroptera, the highest EPT and the largest proportion of EPT taxa. As a result, HBI values were lowest for Hester-Dendy samples because Ephemeroptera tend to be sensitive taxa with low tolerance values. Beyond a single season, we have shown that Hess samples collected more taxa than Hester-Dendy samples across five consecutive years of sampling in the Niobrara River (Tronstad and Hotaling 2017). Dipnets performed consistently poorer than both Hester-Dendy and Hess samples in terms of the number of unique taxa recovered. Similarly, Hester-Dendy samples collected lower taxonomic diversity compared to kicknet samples (McCabe et al. 2012; Letovsky et al. 2012), sweep nets and stovepipe cores (Turner and Trexler 1997) in other aquatic ecosystems. Quantitative samplers (e.g., Surber and Hess samplers) collected similar (Buss and Brges 2008) or more taxa than kicknets (Gillies et al. 2009) and box samplers (O’Connor et al. 2004). In the Niobrara River, Hess samples contained more than twice as many taxa as dipnets at two of the sites. Thus, our study lends additional support to previous findings that quantitative sampling (e.g., Hess or Surber) outperforms other methods by collecting more taxa overall, more unique taxa, and by sampling natural features, a more representative view of the natural community (Tronstad and Hotaling 2017).

Hester-Dendy and Hess samples suggested that invertebrates were fairly evenly distributed in the sampled assemblage based on taxonomic evenness. We calculated taxonomic evenness as Shannon’s diversity index divided by the log_10_ of richness. A value near zero indicates that the assemblage is dominated by a few taxa whereas a value near one indicates that the abundance of each taxon is similar. Mean richness for composited samples were close to one for both Hess and Hester-Dendy samples; however, dipnet samples had a mean value of 0.55, suggesting substantial bias in the assemblage towards high density taxa (Table 1). Specifically, our dipnet samples had a high abundance of Amphipoda. Our results indicated that taxonomic evenness should only be compared to other dipnet samples and dipnets likely underestimate the evenness of the invertebrate community being studied.

We recommend sampling quantitatively (e.g., Hess) for aquatic invertebrate biomonitoring studies when streams are wadeable. In our study, Hess samples collected the most taxa overall, yielded an intermediate HBI value and we expect most closely reflected the natural community because we sampled natural, benthic features in the stream. A stovepipe core would likely produce similar results. For sample processing, we recommend collecting replicate samples in the field, especially when variance is important for detecting changes (e.g., over time or differences among variables of interest). Generally, composite samples lack the statistical power to detect changes in variables of interest. Choosing the most appropriate sampling method paired with processing each replicate individually will provide the most valuable experimental design in most cases, particularly because replicates can always be electronically combined after the fact but the reciprocal is not true.

## Acknowledgments

We thank Katrina Cook, Linda Cooper, Isaac Dority, Heather Hicks, Ariele Johnson, Alexis Lester, Tresize Tronstad and Sarah Wannemuehler for field and laboratory assistance. Robert Manasek and James Hill of the National Park Service provided logistical and field support, as well as the opportunity to work at Agate Fossil Beds National Monument. The project was supported by the National Park Service. Discussions with Brett Marshall were helpful in developing the manuscript.

