## Supplementary Materials for "To composite or replicate: how sampling method and protocol differences alter stream bioassessment metrics"

### To composite or replicate: how bioassessment metrics change depending on sampler type and protocols

**Affiliations:**
[1] Wyoming Natural Diversity Database, University of Wyoming, Laramie, Wyoming
[2] Northern Great Plains Inventory and Monitoring Program, National Park Service, Rapid City, South Dakota
[3] School of Biological Sciences, Washington State University, Pullman, Washington

**Supplemental material 1** Mean abundance (individuals/sample) of invertebrates collected from three sites along the Niobrara River at Agate Fossil Beds National Monument in 2016 using Hester-Dendy samplers. An * marks taxa not found in Hess samples and # marks taxa not found in dipnet samples.

| | East | Middle | Ranch |
|---|---|---|---|
| **Annelida** | | | |
| *G. complanata* # | 1 | | |
| *H. stagnalis* # | | 1 | |
| Oligochaeta | 3 | 3 | 3 |
| **Collembola** | **3** | **1** | **2** |
| **Crustacea** | | | |
| **Amphipoda** | | | |
| Gammaridae | | | |
| *Gammarus* | 14 | 201 | |
| Hyalellidae | | | |
| *Hyalella* | 74 | 175 | 5 |
| **Ostracoda #** | 6 | 10 | |
| **Gastropoda** | | | |
| Ancylidae # | 67 | 23 | 1 |
| Lymnaeidae | 2 | | |
| Physidae # | 3 | 4 | |
| Planorbidae * # | 1 | | |
| **Insecta** | | | |
| **Coleoptera** | | | |
| Dytiscidae | | | |
| *Liodessus* * | 1 | | |
| Gyrinidae | | | |
| *Gyrinus* | 3 | | |
| Haliplidae | | | |
| *Haliplus* # | 1 | | |
| Hydrophilidae * | | | |
| *Enochrus* * # | 1 | | |
| *Hydrochus* * # | 1 | 1 | |
| **Diptera** | | | |
| Ceratopogonidae | | | |
| *Bezzia/Palpomyia* | 1 | 1 | |
| *Culicoides* # | 1 | | 2 |
| Chironomidae | | | |
| Chironomidae (pupae) | 1 | 10 | 1 |
| Non-Tanypodinae | 66 | 187 | 7 |
| Tanypodinae | 4 | 16 | |
| Empididae | | | |

| | | | |
|---|---|---|---|
| *Oreogeton* # | | 1 | |
| Psychodidae * # | | 1 | |
| Simuliidae | | | |
| *Simulium* # | 2 | 56 | 23 |
| **Ephemeroptera** | | | |
| Baetidae | 13 | 9 | 28 |
| Caenidae | | | |
| *Caenis* | 7 | 49 | 30 |
| Heptageniidae | | | |
| *Heptagenia* | 3 | | 31 |
| Leptohyphidae | | | |
| *Tricorythodes* * # | | 1 | |
| Leptophlebiidae | | | |
| *Paraleptophlebia* * # | 8 | 141 | 70 |
| **Hemiptera** | | | |
| Belostomatidae | | | |
| *Belostoma* | | 2 | |
| **Odonata** | | | |
| Aeshnidae | 4 | | |
| Coenagrionidae | | | |
| *Coenagrion/Enallagma* | 6 | 21 | 1 |
| **Trichoptera** | | | |
| Hydropsychidae | | | |
| *Cheumatopsyche* | 1 | 5 | 29 |
| Hydroptilidae | | | |
| *Ochrotrichia* * # | 1 | | |
| *Oxyethira* | 2 | 14 | |
| Polycentropodidae | | | |
| *Polycentropus* # | 1 | | |
| **Turbellaria** | | | |
| *Turbellaria* # | | 15 | |

**Supplemental material 2** Mean density (individuals/sample) of invertebrates collected from three sites along the Niobrara River at Agate Fossil Beds National Monument in 2016 using a Hess sampler. An * marks taxa not found in Hester-Dendy samples and # marks taxa not found in dipnet samples.

| | East | Middle | Ranch |
|---|---|---|---|
| **Annelida** | | | |
| *Erpobdella* * # | | 4 | 2 |
| *G. complanata* # | 3 | 4 | |
| *H. stagnalis* # | | 3 | |
| Oligochaeta | 65 | 52 | 227 |
| **Arachnida** | | | |
| Acari * # | | | 1 |
| **Bivalva** | | | |
| **Bivalvia** | | | |
| Sphaeriidae | 9 | | |
| **Collembola** | **2** | **4** | **5** |
| **Crustacea** | | | |
| **Amphipoda** | | | |
| Gammaridae | | | |
| *Gammarus* | 12 | 42 | 6 |
| Hyalellidae | | | |
| *Hyalella* | 175 | 124 | 8 |
| **Decapoda** | | | |
| Cambaridae | | | |
| *O. neglectus* * | 1 | 3 | 6 |
| **Ostracoda** # | 1 | 4 | |
| **Gastropoda** | | | |
| Ancylidae # | 6 | 4 | 4 |
| Lymnaeidae | 1 | | 2 |
| Physidae # | 3 | 2 | |
| Spheariidae | 9 | | |
| **Insecta** | | | |
| **Coleoptera** | | | |
| Dytiscidae | | | |
| *Agabetes* * # | 1 | | |
| *Celina* * # | | 1 | |
| *Hydrovatus* * # | | 1 | |
| *Ilybius* * # | 1 | | |
| Lampyridae * # | | 1 | |
| Gyrinidae | | | |
| *Gyrinus* | 7 | 2 | |
| Haliplidae | | | |

| | | | |
|---|---|---|---|
| *Haliplus #* | | 1 | |
| *Peltodytes * #* | | 1 | |
| Helophoridae | | | |
| *Helophorus * #* | 1 | 1 | |
| **Diptera** | | | |
| Ceratopogonidae | | | |
| *Bezzia/Palpomyia* | 5 | 2 | |
| *Ceratopogon #* | 5 | 2 | |
| *Culicoides* | 15 | | 3 |
| *Probezzia * #* | | 2 | |
| Chironomidae | | | |
| Chironomidae (pupae) | 39 | 6 | 2 |
| Non-Tanypodinae | 68 | 409 | 26 |
| Tanypodinae | 7 | 5 | |
| Dixidae | | | |
| *Dixa * #* | | 1 | |
| Empididae | | | |
| *Oreogeton * #* | 5 | | |
| Ephydridae | | | |
| *Notiphila * #* | 13 | 5 | |
| Simuliidae | | | |
| *Simulium* | 32 | 6 | 5 |
| Stratiomyiidae | | | |
| *Odontomyia ** | 1 | 1 | |
| *Stratiomys * #* | | 1 | |
| Tabanidae * # | 3 | 7 | 1 |
| Tipulidae | | | |
| *Dicranota * #* | | 2 | |
| *Ormosia * #* | 11 | | |
| **Ephemeroptera** | | | |
| Baetidae | 3 | 8 | 6 |
| Caenidae | | | |
| *Caenis* | 7 | 156 | |
| Ephemeridae | | | |
| *Hexagenia * #* | | | 3 |
| Heptageniidae | | | |
| *Heptagenia* | 1 | | 4 |
| **Hemiptera** | | | |
| Belostomatidae | | | |
| *Belostoma* | 1 | 2 | |
| Corixidae | | | |
| *Hesperocorixa * #* | | | 1 |
| Gerridae | | | |

| | | | |
|---|---|---|---|
| *Gerris * #* | | 1 | |
| Notonectidae | | | |
| *Notonecta ** | | 1 | |
| **Odonata** | | | |
| Aeshnidae | | | |
| *Anax ** | | 2 | |
| Coenagrionidae | | | |
| *Coenagrion/Enallagma* | 10 | 76 | |
| Gomphidae * # | 1 | 1 | |
| Libellulidae * | | 3 | |
| **Trichoptera** | | | |
| Hydropsychidae | | | |
| *Cheumatopsyche* | 1 | | 8 |
| Hydroptilidae | | | |
| *Oxyethira* | | 7 | |
| Polycentropodidae | | | |
| *Polycentropus #* | | 1 | |
| **Nematoda *** | **2** | | |
| **Nematomorpha * #** | **6** | **5** | **1** |
| **Turbellaria #** | | **3** | |

**Supplemental material 3** Abundance (count) of invertebrates collected from three sites along the Niobrara River at Agate Fossil Beds National Monument in 2016 using a D-frame dipnet. An * marks taxa not found in Hess samples and # marks taxa not found in Hester-Dendy samples.

| | East | Middle | Ranch |
|---|---|---|---|
| **Annelida** | | | |
| Oligochaeta | | 5 | 15 |
| **Bivalva** | | | |
| Sphaeriidae | | | 2 |
| **Collembola** | | | **16** |
| **Crustacea** | | | |
| **Amphipoda** | | | |
| Gammaridae | | | |
| *Gammarus* | 16 | 28 | 3 |
| Hyalellidae | | | |
| *Hyalella* | 509 | 134 | 34 |
| **Copepoda** | | | |
| Harpacticoida * # | | | 4 |
| **Decapoda** | | | |
| Cambaridae | | | |
| *O. neglectus #* | 4 | 3 | |
| **Gastropoda** | | | |
| Lymnaeidae | | | 5 |
| Spheariidae # | | | **2** |
| **Insecta** | | | |
| **Coleoptera** | | | |
| Dytiscidae | | | |
| *Agabus * #* | 1 | | |
| *Coptotomus * #* | | 1 | |
| *Liodessus ** | 1 | | |
| Gyrinidae | | | |
| *Gyrinus* | 6 | 1 | |
| Hydrophilidae* | | | |
| *Hydrophilidae* | | | 1 |
| *Tropisternus * #* | | 2 | |
| **Diptera** | | | |
| Ceratopogonidae | | | |
| *Bezzia/Palpomyia* | | 2 | |
| *Ceratopogon #* | | | 96 |
| Chironomidae | | | |
| Chironomidae (pupae) | | 2 | 1 |
| Non-Tanypodinae | 12 | 223 | 63 |
| Tanypodinae | 3 | 12 | |

| | | | |
|---|---|---|---|
| Simuliidae | 17 | 18 | 62 |
| Stratiomyiidae | | | |
| *Odontomyia #* | | | 1 |
| Tipulidae | | | |
| *Tipula * #* | | 1 | |
| **Ephemeroptera** | | | |
| Baetidae | 2 | 3 | 37 |
| Caenidae | | | |
| *Caenis* | 7 | 41 | 5 |
| Ephemeridae | | | |
| *Ephemera * #* | | | 9 |
| Heptageniidae | | | |
| *Heptagenia* | | | 39 |
| **Hemiptera** | | | |
| Belostomatidae | | | |
| *Belostoma* | | 2 | |
| Notonectidae | | | |
| *Notonecta #* | | 1 | |
| **Odonata** | | | |
| Aeshnidae | | | |
| *Anax #* | | 1 | 1 |
| Calopterygidae | | | |
| *Calopteryx * #* | | | 1 |
| *Hetaerina * #* | 1 | | |
| Coenagrionidae | 10 | | |
| *Coenagrion/Enallagma* | | 49 | |
| Libellulidae | | | |
| *Libellula #* | | 3 | |
| **Trichoptera** | | | |
| Hydropsychidae | | | |
| *Cheumatopsyche* | | | 3 |
| Hydroptilidae | | | |
| *Oxyethira* | | 5 | 1 |
| **Nematoda #** | | | **2** |
